# Rare genetic variation and balanced polymorphisms are important for survival in global change conditions

**DOI:** 10.1101/422261

**Authors:** Reid S. Brennan, April D. Garrett, Kaitlin E. Huber, Heidi Hargarten, Melissa H. Pespeni

## Abstract

Standing genetic variation is important for population persistence in extreme environmental conditions. While some species may have the capacity to adapt to predicted average future global change conditions, the ability to survive extreme events is largely unknown. We used single generation selection experiments on hundreds of thousands of *Strongylocentrotus purpuratus* sea urchin larvae generated from wild-caught adults to identify adaptive genetic variation responsive to moderate (pH 8.0) and extreme (pH 7.5) low pH conditions. Sequencing genomic DNA from pools of larvae, we identified consistent changes in allele frequencies across replicate cultures of both conditions and observed increased linkage disequilibrium around selected loci, revealing selection on recombined standing genetic variation. We found that loci responding uniquely to either selection regime were at low starting allele frequencies while variants that responded to both pH conditions (11.6% of selected variants) started at high frequencies. Loci under selection performed functions related to energetics, pH tolerance, cell growth, and actin/cytoskeleton dynamics. These results highlight that persistence in future conditions will require two classes of genetic variation: common, pH-responsive variants maintained by balancing selection in a heterogeneous environment, and rare variants, particularly for extreme conditions, that must be maintained by large population sizes.

## Introduction

As temperatures increase and oceans become more acidic, many marine species are at risk of decline and extinction [1]. In addition to changing global averages, the frequency and intensity of extreme events are increasing [2]. While organisms may acclimate through physiological plasticity or migrate to suitable habitats, some amount of genetic adaptation will be necessary for population persistence, particularly in the context of extreme events [3]. Indeed, extreme events rather than average conditions play an important role in setting physiological and biogeographic limits of species [4-6]; however, the genetic mechanisms that allow rapid adaptation to extreme conditions have rarely been explored [7].

Evolve or select and resequence, a type of experimental evolution, is a powerful and promising approach for understanding the capacity for adaptation to future conditions and extreme events. Selection can be artificially induced to directly measure adaptive genetic responses [8-11]. While these studies typically leverage 15-20 generations of selection [10,12], empirical studies have found that adaptive genetic variation can be identified with a single generation of selection by maximizing replication and the number of outbred individuals or offspring used [13-15], making this approach applicable to a broad diversity of systems.

The purple sea urchin, *Strongylocentrotus purpuratus*, is an ideal model for understanding the process of adaptation from standing genetic variation. This species is found in inter-and subtidal rocky reefs and kelp forests from Alaska to Baja California, Mexico in the California Current Marine Ecosystem (CCME). They experience high heterogeneity in temperature and pH conditions in both time and space [16-19], increasing the likelihood of the presence of adaptive standing genetic variation [20-22]. Large census and effective population sizes contribute to high standing genetic variation [23,24] and there is evidence of adaptation to local temperature and pH conditions [25]. Previous studies have shown that *S. purpuratus* larvae have the physiological and genetic capacity to adaptively respond to an acidifying ocean [15,18,26], but these studies have focused on relatively mild low pH conditions and have not investigated the capacity to respond to extreme low pH conditions that are predicted to increase in frequency in the near future [2,17].

Here, we performed single generation selection experiments in moderate (pH 8.0) and extreme low (pH 7.5) pH conditions using purple sea urchin larvae generated from 25 wild-caught adults. We used pooled sequencing of genomic DNA to test the hypotheses that (1) larvae before and after treatments will show consistent changes in allele frequency across replicate cultures due to selection for genetic variation that impacts survival in low pH conditions; (2) there will be unique genetic variation responsive to extreme versus moderate low pH conditions; (3) genomic patterns of variation, such as linkage disequilibrium (LD) and starting allele frequencies, will provide insight into the evolutionary mechanisms that maintain adaptive standing genetic variation for survival in future global change conditions.

## Methods

### Sample collection & experiment

Purple sea urchin adults (San Diego, CA, USA; n=25 total: 14 females and 11 males) were induced to spawn using 0.5M KCl in Instant Ocean artificial seawater (ASW) (Instant Ocean, Blacksburg, VA) at 14°C and salinity of 35ppt. For each female, 200,000 eggs were placed in each treatment (moderate (pH 8.0) and extreme (pH 7.5) low pH) and fertilized by evenly pooled sperm from all males. We chose these pH conditions based on empirical data; the current average open ocean and intertidal pH is 8.1, though conditions in the CCME frequently drop to pH 8.0 (daily), and rarely drop as low as pH 7.5 (once in a three month upwelling period; [16,17,19,27]). Fertilized eggs were pooled across all females by pH and seeded into four replicate culturing vessels per treatment (37,000 eggs per 3.7 L vessel). Developing embryos were sampled at day 1 and day 7 post fertilization for morphometric, mortality, and genomic analyses. See supplemental methods for expanded details and Table S1 for water chemistry.

### Morphometrics

Seven-day old 4-armed pluteus larvae were photographed for morphometric analysis using a Photometrics Scientific CoolSNAP EZ camera (Tuscon, AZ) connected to a Zeiss Axioscop 2 compound microscope (Jena, Germany). Larval body size data were analyzed in R [28] with a generalized linear mixed model in the package lme4 [29], with pH as a fixed effect and culturing vessel as a random effect.

### DNA sequencing, Mapping, & SNP-calling

DNA was extracted from pools of approximately 15,000 to 20,000 surviving larvae from each replicate vessel using a Zymo ZR-Duet DNA/RNA MiniPrep Plus Kit (Zymo, Irvine, CA). DNA was shipped to Rapid Genomics (Gainesville, FL) for library prep, capture, and sequencing. DNA was captured with 46,316 custom probes designed to capture two 120 base pair regions per gene: one within exon boundaries and one in putative regulatory regions, within 1000 bp upstream of the transcription start site. Barcoded samples were pooled and sequenced using 150 bp paired-end reads on one lane of an Illumina HiSeqX.

Raw paired-end reads were quality trimmed and mapped to the *S. purpuratus* genome 3.1 (build 7) with bwa mem [30]. Variants were identified using *mpileup* in Samtools and filtered for minor allele frequency (maf) of 0.01, quality greater than 20, bi-allelic SNPs only, and no missing data. We used only variants where each pool was sequenced to a depth of > 40x and with an average minimum depth across all pools > 50x, as recommended for pooled sequencing by Schlotterer et al. [31]. Mean maximum coverage cutoff was 372. Finally, we removed off target variants, any variant greater than 2kb from a probe. This filtering resulted in 75,368 variant sites. See supplemental methods for more details. Code for data processing and analyses can be found on GitHub: https://github.com/PespeniLab/urchin_single_gen_selection.

### Detecting changes in allele frequency

A principal components analysis (PCA) was used to visualize genome-wide relationships between sample allele frequencies using the R package *pcadapt* version 4.0 [32]. We calculated nucleotide diversity for each treatment using a sliding window approach in Popoolation with the *variance-sliding* command [33]. Window size was set to 400 bp with step size of 200 bp.

Loci under selection should increase in frequency consistently among replicates due to differential survival of larvae with adaptive genotypes. Patterns of genetic variation following a selective sweep are typically observed after many generations of selection. However, over a single generation of selection, using wild-caught, outbred parents and 37,000 offspring with recombined genomes, adaptive alleles can be present on multiple different genomic backgrounds or haplotypes and selective mortality can result in a signal of elevated LD that decays with physical distance. In this experiment, selective mortality is analogous to a soft sweep where multiple individuals or haplotypes (from standing variation) carry the adaptive allele that is under selection [33]. The consistency of allele frequency changes and increases in LD among replicates can be used to distinguish selected loci from those undergoing neutral drift.

Cochran-Mantel-Haenszel (CMH) tests, a method for identifying consistent changes in allele frequency in evolution experiments [10], were conducted in R (*mantelhaen.test*) comparing allele frequency estimates from the four replicates at time point zero (T_0_, day 1, pH 8.0) to the four replicate vessels at day 7, pH 8.0 and to the four replicates at day 7, pH 7.5. Significance thresholds were determined using a Bonferroni correction. We considered any SNPs within the same gene to be non-independent due to linkage and calculated the number of tests as the number of independent gene regions; this resulted in a threshold of 0.05/9828 = 5.09e-6. This is an alternative to the overly conservative Bonferroni correction that ignores linkage and considers every site independent [34], and is more stringent than a FDR approach with these data (*P* < 0.004).

We compared LD estimates among pairs of SNPs across the genome for selected (CMH *P* < 5.09e-6) and neutral (non-selected, CMH *P* ≥ 5.09e-6) variants using LDx [35] (supplemental methods). LDx uses a maximum likelihood approach and leverages haplotype information from single reads to estimate LD between pairs of variants over short distance. To estimate the decay in LD, we regressed the log of the physical distance with LD between base pairs for each treatment group with replicate as a random effect using the R package *nlme* [36]. To assess whether the levels of LD present around selected sites could be due to the low number of variants relative to neutral sites, we randomly subsampled all variants 500 times to match the number of selected loci for both pH 7.5 and 8.0. We compared the observed slope of decay and the intercept to permuted distributions.

We used folded allele frequencies to assess if the distribution of starting frequencies of adaptive loci were distinct from the distribution of neutral loci. To ensure the patterns observed were not due to bias of the CMH statistic towards detection of variants at low starting allele frequencies, the observed distributions were compared to permuted distributions with no true biological signal as described in the supplemental methods.

Gene ontology (GO) enrichment was performed using the weight algorithm in topGO version 2.22.0 [37]. GO terms for each gene were retrieved from EchinoBase (http://www.echinobase.org/). Enrichment tests were conducted for each set of significant variants and genes that had any SNPs in genic or regulatory regions were considered the target set. Any gene with multiple significant variants was only considered once.

## Results

### Strength of selection

Mean total larval body length was lower in pH 7.5 (mean ± standard error: 239.1 ± 3.1μm) as compared to pH 8.0 (320.9 ± 2.9 μm) (Fig. 1, *P* < 0.001). Mortality estimates (the proportion of larvae dead at day 7, supplemental methods) differed in the same direction as body length, though not significantly, where pH 7.5 had a higher mean mortality of 51.5% + 11.3% and pH 8.0 was 50.8% + 5.4% (*P* = 0.89). The median selection coefficient (supplemental methods), was 0.210 and 0.206 for pH 7.5 and 8.0 selected loci, respectively, and was marginally different (*P* = 0.07). These results suggest that physiological and selective impact was stronger in pH 7.5 than pH 8.0.

**Figure 1:**
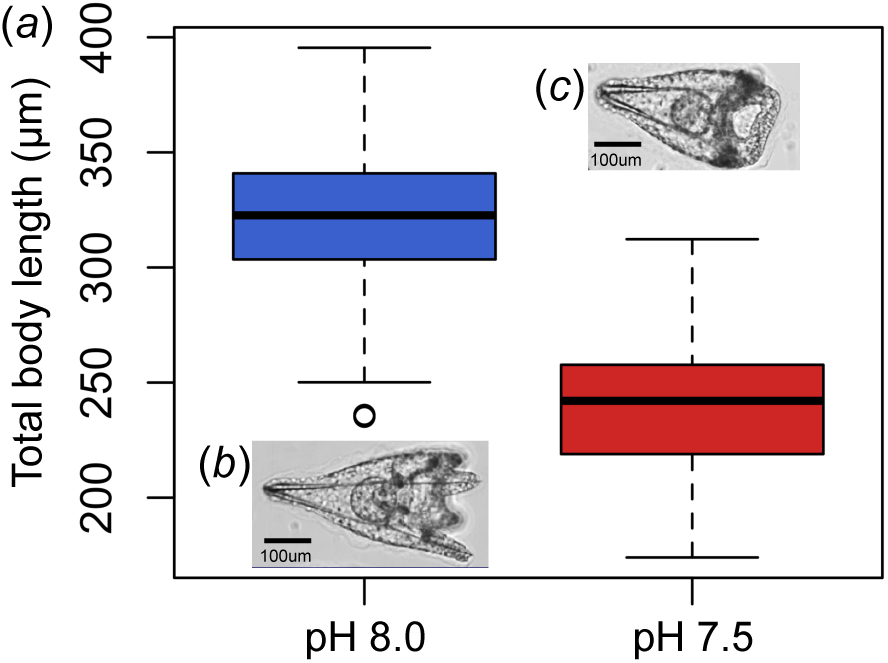
Morphometric analysis of total larval body length. (a) Boxplot of body size in each selection regime. Inset pictures show representative individuals found in pH 8.0 (b) and 7.5 (c).

### Consistent allele frequency changes among replicate selection lines

Across the twelve pools of larvae (four replicates of each: T_0_ and after 7 days in pH 8.0 and pH 7.5), we identified 75,368 variable sites present in or near 9,828 genes with an average coverage of 123x per site per sample. PCA showed that samples clustered by day and treatment (Fig. 2a), indicating consistent allele frequency estimates from each biological replicate and consistent changes in allele frequency in response to treatment conditions. pH 7.5 samples showed the largest shift from the starting allele frequencies as expected with increased selective mortality due to treatment (Fig. 2a). Note that one of the D7 pH 8.0 samples was an outlier. To ensure that this sample did not reduce power, we removed this sample, down sampled all replicates to n=3, and reran analyses; results were consistent with and without this sample.

**Figure 2:**
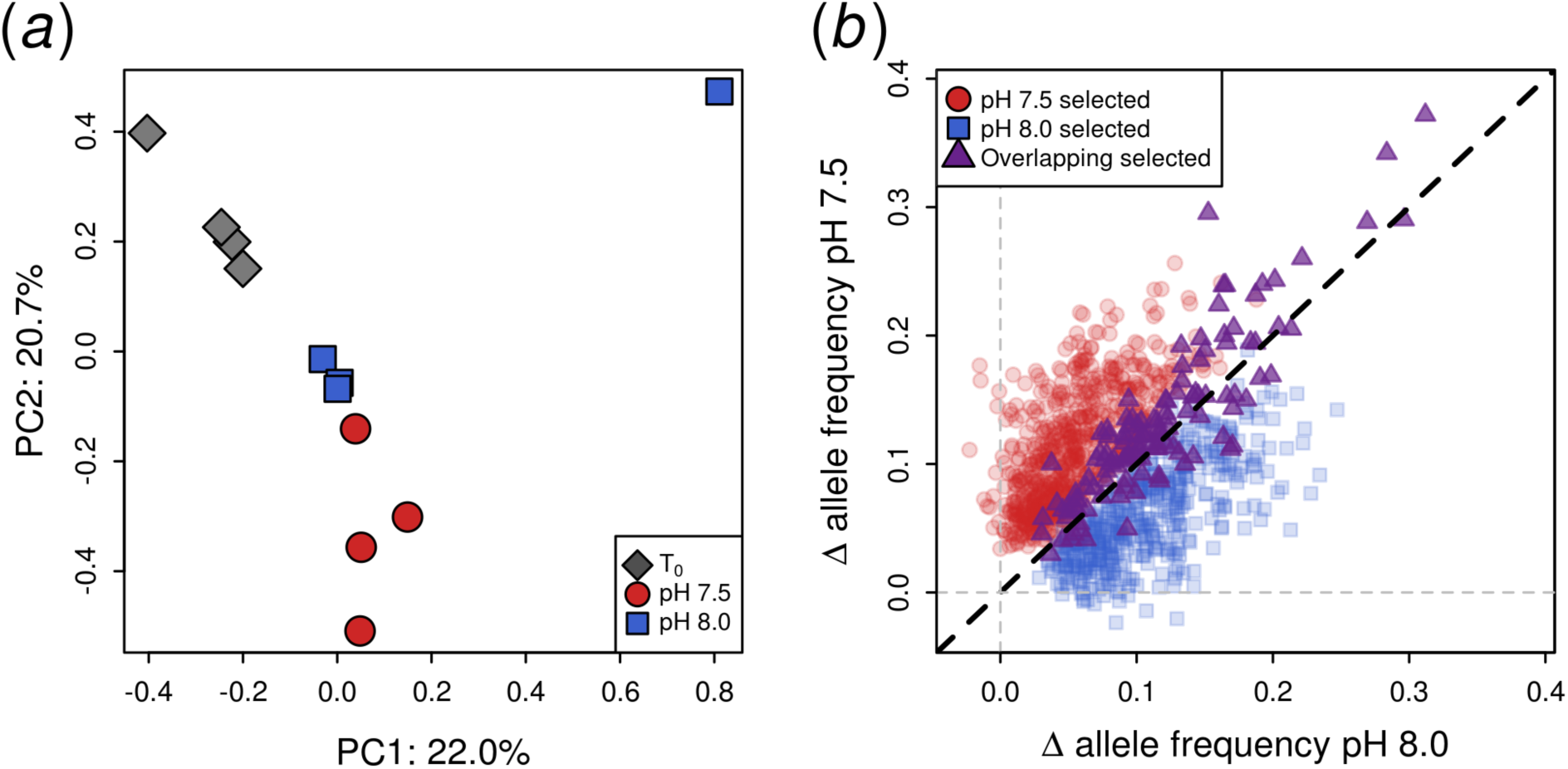
(a) Principal component analysis of allele frequencies for all 75,368 SNPs at time point zero (T_0_) and after seven days in the two pH treatments. (b) Mean change in allele frequency for variants showing significant shifts under pH 7.5 and 8.0, and to both pH regimes. Dashed black line represents the 1:1 expectation.

We found 787 variants (in or near 606 genes) with consistent changes in allele frequency in response to selection at pH 7.5 and 572 variants (470 genes) that changed in response to pH 8.0 (CMH, *P*<5.09e-06; Fig. 2b; see Table S2 for allele frequency data and CMH results). The mean change in allele frequency was 0.115 ± 0.049 (mean ± SD) and 0.110 ± 0.046 for pH 7.5 and pH 8.0 responsive variants, respectively, while the genome-wide mean change was 0.028 ± 0.030. 128 (11.6%) of pH 7.5-and pH 8.0-responsive variants overlapped (100 genes).

Of the total assayed variants, 50,691 (67%) were in genic regions while 24,677 (33%) were in regulatory regions. Matching expectations of chance sampling, for pH 7.5 selected loci, 537 (68%) were genic and 250 (32%) were regulatory (chi-squared, *P* > 0.05). Similarly, pH 8.0 selected loci consist of 401 (70%) genic and 171 (30%) regulatory loci (chi-squared, *P* > 0.05). Given the rapid decay in linkage disequilibrium (see below), these results suggest that there are both important coding and putative regulatory pH-responsive loci segregating in populations.

### Signals of shared and pH-specific selection

Shifts in allele frequency showed unique and shared signals between pH selection regimes (Fig. 2b, S1). Significant loci specific to pH 7.5 show a correlation in average allele frequency change of 0.67 with their non-significant pH 8.0 counterparts (*P* < 0.001) suggesting that the same loci were responding to treatment though to a lesser degree in pH 8.0; pH 8.0 significant loci reveal the same pattern with a correlation of 0.71 (*P* < 0.001). As expected, loci that were identified as significant in both treatments showed the strongest correlation in allele frequency change (0.89, *P* < 0.001). Interestingly, the loci with the most extreme shifts in allele frequency were significant in both selection regimes (upper right quadrant of Fig. 2b, Fig. S2). In comparison, the correlation of allele frequency change in non-significant loci was 0.19.

### Elevated LD around selected loci

Selected loci in both pH treatments showed strong linkage disequilibrium with the slowest decay in LD for SNP pairs involving pH 7.5 selected loci (estimated slope of regression of log distance and LD ± standard error: −0.069 ± 0.005), followed by pH 8.0 selected loci (slope: −0.072 ± 0.006); both of these decay rates were slower than the −0.111 mean from the permutations and fell outside of the 95% distribution (pH 7.5 slope: −0.134 to −0.085; pH 8.0 slope: −0.145 to −0.079; Fig. 3a). We observed rapid decay of LD within 100 base pairs, which is expected given the high levels of genetic diversity, large effective population size, high fecundity, and high gene flow of this species. As expected, the observed intercepts, LD at zero distance, (0.306 ± 0.008 and 0.283 ± 0.009) for pH 7.5 and 8.0 selected loci were not significantly different from neutral expectations as both sets fall within the range of permutation 95% distributions (pH 7.5: 0.310 (0.298, 0.382); pH 8.0: 0.283 (0.282, 0.393)).

**Figure 3:**
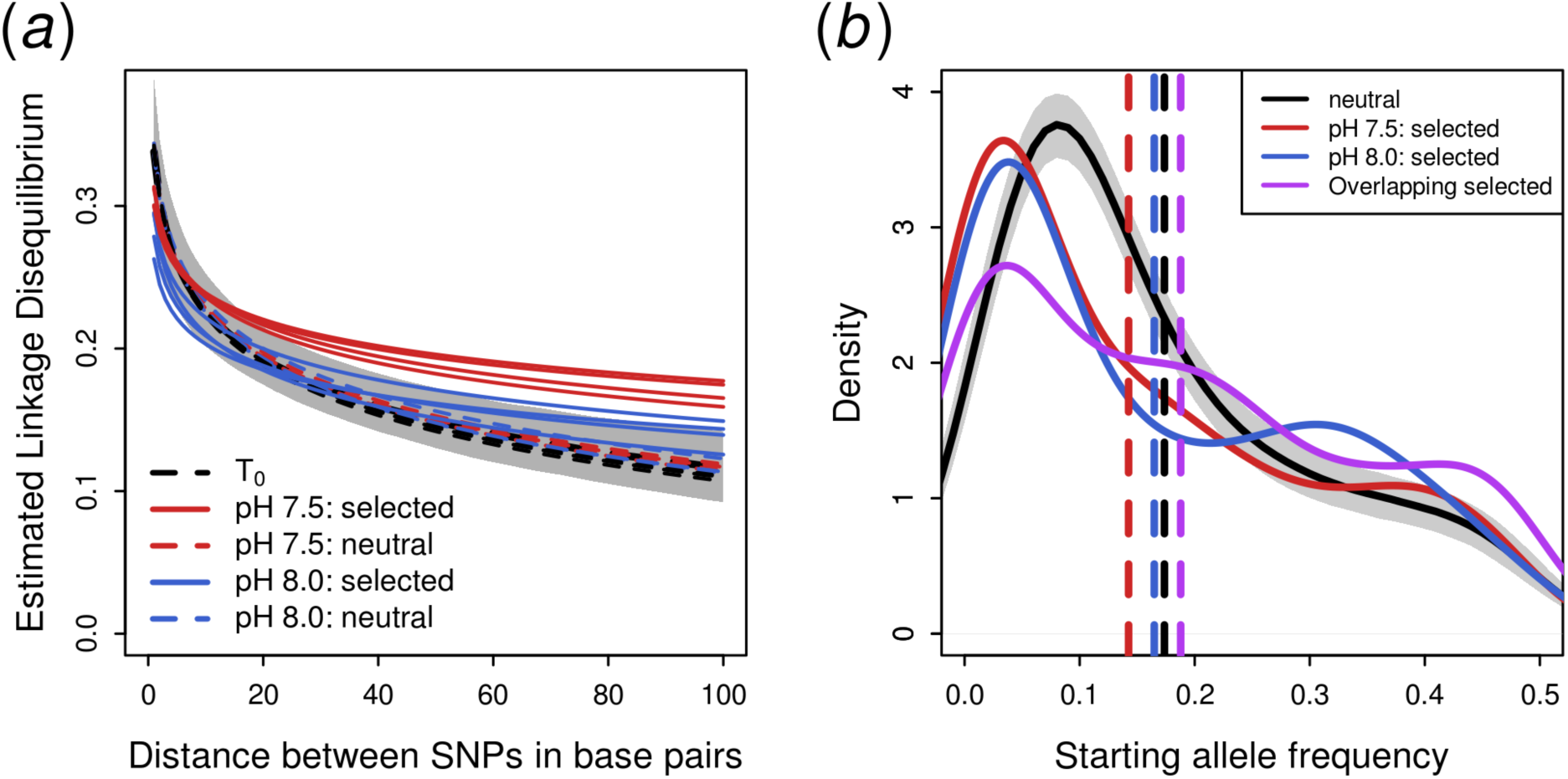
(a) Decay in linkage disequilibrium with physical distance between SNP pairs. Dashed lines represent decay around non-significant, putatively neutral loci and solid red and blue lines represent decay around pH 7.5 and 8.0 putatively selected variants, respectively. Solid grey shading is the 95% distribution of 500 permutations of down sampled random variants across the genome (based on pH 8.0 for visualization). Note that each of the four replicates per treatment are plotted though may be difficult to see due to the highly concordant decay. (b) Folded allele frequency at T_0_. Colored solid lines are the density plot distributions for each set of selected loci. The solid black line represents the median expectation of neutral loci from 1000 permutations of down sampling to the same number of variants as selected loci; grey shading is the 95% distribution of the permutation. Dashed vertical lines show the mean allele frequencies of each set of variants.

### Maintenance of pH-adaptive alleles in natural populations

To understand how variants are segregating and maintained in populations, we explored the starting allele frequencies (T_0_) for loci identified as responsive to pH 7.5 (mean ± standard error: 0.15 ± 0.01), pH 8.0 (0.16 ± 0.01), and to both treatments (0.19 ± 0.01; Fig. 3b). The distributions of starting allele frequencies for pH 8.0 and 7.5 selected loci were significantly lower than neutral loci (0.17 ± 0.0004; two-sided Kolmogorov–Smirnov tests, *P* < 0.001), while the overlapping selected loci were shifted towards higher frequencies (*P* < 0.001). Loci responsive to the most extreme selection regime, pH 7.5, were lower in starting allele frequency than loci responsive to both pH treatments (*P* = 0.03) and to pH 8.0 alone, though this effect was not significant for the latter (*P* = 0.08). Conversely, loci responsive to only pH 8.0 did not have lower starting allele frequency than those responding to both treatments (*P* = 0.1). These patterns hold when selected loci were polarized by the allele increasing in frequency in response to pH selection (the putative adaptive allele; Fig. S3). We also compared observed allele frequency distributions of selected loci to distributions of the same number of randomly sampled loci across the genome (Fig. 3b). The distributions for pH 7.5, pH 8.0, and overlapping selected variants were significantly different from all 1000 permutations (*P* < 0.001). Lastly, permutations showed that the CMH statistic was not biased towards loci that started at low frequencies; randomly shuffled sample IDs were not enriched for low starting allele frequencies (Fig. S4). Finally, we found that loci with higher minor allele frequencies showed the greatest changes in frequency in response to selection (Fig. S5), which matches theoretical and simulation expectations [8]. The excess of low allele frequency variants among selected loci is consistent across functional classes of variants (non-synonymous, synonymous, intronic, regulatory; Fig. S6).

Genetic diversity (π) was 0.0158 at T_0_ and decreased to 0.0129 and 0.0128 after seven days at pH 8.0 and 7.5, respectively (one-sided Wilcoxon rank-sum *P* < 0.001). The decrease in diversity between the pH treatments was not different (*P* = 0.12), demonstrating that, while the selection regimes were of different magnitude, the relative loss of genetic diversity through time was similar.

Functional enrichment analysis revealed that genes under selection were enriched for specific biological functions (Table S3). We observed enrichment for 17, 10, and 14 biological processes GO terms for pH 7.5, pH 8.0, and overlapping variants, respectively. Of these, only hexose metabolic process (GO:0019318) was significant across all three variant sets. Other related terms shared across sets included those involved in cell growth (GO:0030307; GO:0032467), metabolism and energy production (GO:0001678; GO:0009267; GO:0019318; GO:0006096; GO:0006869; GO:0009061). pH 7.5 was enriched for unique functional processes, particularly those involved in cytoskeleton remodeling (GO:0051639; GO:0008154) and intracellular pH regulation (GO:0007035). pH 8.0 and overlapping variants showed limited signal of unique enrichment, see Table S3 for all enriched categories.

## Discussion

We show that genetic response to moderate and extreme low pH relies on shared and unique mechanisms and that rare variants are important for survival in low pH conditions. We also identify a class of variants responsive to both pH levels (11.6% of pH-responsive loci) that are maintained at higher frequencies in natural populations, potentially by balancing selection in a spatially and temporally variable pH environment. We further demonstrate the utility of single generation selection experiments to identify the genetic targets of adaptation, which is particularly useful for testing capacity for response to conditions that will be experienced more frequently in the future but are rare events in nature today. Using sequence capture of genomic DNA, we quantified shifts in allele frequency that represent differential survival of genotypes during low pH selection in sea urchins. This approach has wide potential application for identifying responsive genetic variants from wild populations. Our results indicate that standing genetic variation maintained with large population sizes will be critical for survival in future environmental conditions.

### Detecting adaptive loci from standing genetic variation

Recent selective sweeps result in decreased genetic variation and increased linkage disequilibrium (LD) around adaptive loci [38,39]. Even in a single generation of selection, we have the potential to detect selective sweeps as increased LD around adaptive loci from standing variation. In the present experiment, adaptive genetic variation can be present in many different genomic backgrounds through: 1) the use of 25 wild-caught, outbred parents, 2) the recombination that occurred between the diploid genomes of each parent during gametogenesis, and 3) experimental fertilization of eggs from each female with sperm from each male. Seeding each replicate with 37,000 embryos further amplified the number of uniquely recombined genomes. We expect that differential survival of larvae with specific combinations of alleles will yield a pattern consistent with soft selective sweeps where adaptive loci are present on multiple haplotypes [21,40,41]; we observe increased LD in regions surrounding putatively adaptive loci to match this expectation. Permutation tests demonstrate that this increase in LD was not a byproduct of our test statistic or sampling noise (Fig. 3a). In addition, the consistent results among replicates for each treatment provide strong evidence that our approach identifies genomic regions that are true targets of selection, rather than due to drift or chance changes in allele frequency.

Typical evolve and resequence studies rely on multiple generations of selection to identify adaptive shifts in allele frequency. To increase power to distinguish between drift and selection, one can increase the selection coefficient, number of generations, population size, number of replicates, or starting genetic diversity [12,42,43]. Because purple sea urchin take two years to reach reproductive maturity [44] we can use only a single generation of selection, but their high fecundity and small offspring size enable us to maximize population size and genetic diversity. This tradeoff is particularly useful for performing selection experiments in long-lived organisms. One starts with outbred wild-caught individuals, which will increase starting genetic diversity relative to inbred lab strains or isofemale lines [12,45,46]. Inference is limited to the genetic diversity present in the starting individuals and may miss variation present at different points in space or time and will miss the potential of new beneficial mutations, which may be an important mode for adaptation [21]. However, starting with outbred parents maximizes the amount of recombined genetic variation, avoiding the large linkage blocks that can plague experimental evolution studies from lab lines. These narrow linkage windows also improve the chances of identifying the specific loci targeted by selection.

### Maintenance of adaptive standing genetic variation

Variants that responded to low pH treatments were present in the starting population (T_0_) at low allele frequencies relative to neutral alleles (Fig. 3b). Low starting allele frequencies of responsive variants could suggest that surviving offspring were from a single parent. However, this is unlikely due to the decay of LD around selected loci, indicating that many different genetic backgrounds were targeted by selection. In addition, the large number of responsive loci and genes suggests that survival in low pH conditions is a polygenic trait, a characteristic of adaptation to complex environments in other experimental evolution systems [47]. Thus it is unlikely that surviving offspring inherited all the adaptive alleles from a single parent.

The low starting allele frequency of uniquely response low-pH adaptive variants argues that these loci are not maintained by balancing selection or overdominance, but by one of three processes. First, the alleles may be beneficial under low pH but neutral or slightly deleterious at ambient conditions, known as conditional neutrality [48]. Alternatively, antagonistic pleiotropy can alter the rank fitness of alleles across different environments, maintaining genetic variation across space or time [49]. Finally, purple sea urchins inhabit rocky reefs across a broad geographic range [24,50] where pH conditions vary due to natural processes such as upwelling [17]. Low pH-responsive loci in the lab have been shown to have allele frequencies correlated with local pH conditions in the field, suggesting local adaptation [25]. High gene flow across the geographic range and migration-selection balance can result in the maintenance of low frequency alleles that are not adaptive in a local environment [51]. Similar selection on low frequency alleles has been observed in *D. melanogaster* during experimental evolution to high temperature [52]. Interestingly, this shift was observed for adaptation to high but not low temperature. Conversely, Barghi et al., find that experimental adaptation to temperature in *Drosophila* recruits alleles at higher frequency than neutral loci [47], providing additional evidence that the CMH statistic is not inherently biased towards low starting minor allele frequencies.

In contrast, loci overlapping in response to both moderate and extreme pH had higher starting allele frequencies than neutral loci and loci responsive to either pH regime alone (Fig. 3b). These “low-pH-essential” alleles may be maintained at higher allele frequency due to balancing selection in the wild. That is, the spatial and temporal heterogeneity of pH conditions experienced by purple sea urchins across the species range and across life history stages likely maintains these alleles at intermediate frequencies through fluctuating selection or spatially balancing selection [53]. We note that these shared loci may also include variants involved in selection for the general lab culture conditions. While the design of this study precludes the isolation of lab-adapted variants, the functional classes of genes we identify here are also responsive to moderate low pH conditions (pH 7.8) in purple sea urchin larvae generated from multiple populations [14,23] and are correlated with environmental pH in wild populations [23]. In addition, lower pH results in more extreme patterns of reduced LD decay, allele frequency change, and morphological impacts than moderate pH conditions, suggesting that, while lab selection may be present, pH selection is the main driver of the results in the present study.

### Mechanisms for response to low pH conditions

Selected loci are enriched for a range of biological processes related to maintaining cellular homeostasis. Across all sets of selected variants, we observe enrichment in processes related to metabolism and energy production (GO:0001678, GO:0009267, GO:0019318, GO:0006096, GO:0006869, GO:0009061). Alterations to metabolic processes, energy demands, and resource allocations are primary responses to low pH environments [54,55]. Previous transcriptomic studies in *S. purpuratus* have shown that these classes of genes are differentially regulated in response to low pH stress [18,56]. The energy required for acid-base regulation is increased under acidic conditions because the calcification process, production of calcium carbonate (CaCO_3_), generates excess protons that must be removed to maintain acid-base balance [55]. We also see shared enrichment for genes involved in cell proliferation and damage control, including those implicated in cell growth and replication (GO:0030307, GO:0008361, GO:0032467) and protein transport (GO:0006606). These results match our morphometric findings of stunted larval growth and suggest that fine control of cellular growth may be important for survival in low pH conditions.

Two genes of interest that showed changes in allele frequency greater than 20% in both pH treatments perform functions related to cytoskeletal and membrane dynamics: ELMOD2 (SP-ELMOD2/SPU_007564), and Focadhesin (KIAA1797/SPU_015184). ELMOD2 had the greatest changes in allele frequency in both treatments and plays an important role in regulating membrane traffic and secretion, phospholipid metabolism, and actin/cytoskeleton dynamics [57-59]. The Focadhesin protein plays an important role in integrating intra-and extracellular information as a subcellular structural protein that receives biomechanical and biochemical signals between the cytoskeleton and the extracellular matrix [60,61]. We identify three SNPs, two of which change amino acids, which show shifts in allele frequency greater than 20% in this putatively low-pH adaptive gene.

Selection in pH 7.5, the edge of what *S. purpuratus* urchins experience in nature, revealed unique adaptive targets to extreme pH selection. We observe enrichment for ‘actin polymerization or depolymerization’ (GO:0008154) and actin filament network formation (GO:0051639). It has been hypothesized that changes in actin abundance are related to cytoskeleton remodeling due to intracellular stress during acclimation of oysters to climate change conditions [62,63]. Evans et al., [16] find enrichment for expression of genes involved in actin folding in *S. purpuratus*, suggesting that the cytoskeleton is a target of pH stress. We also found enrichment for vacuolar acidification (GO:0007035); regulation of pH in intracellular compartments can maintain pH homeostasis when extracellular pH is altered [64].

We identify putative adaptive alleles in proteins that play important roles in bicarbonate uptake and pH regulation. TASK2 is a pH-sensitive K+ transporter [65] and is an important component of bicarbonate (HCO3-) uptake in higher eukaryotes [66]. We identify pH 7.5-selected alleles in TASK2 (SPU_003613; Fig. 4). Recent studies in purple sea urchin larvae have shown that the regulation of intracellular pH is linked to biomineralization and that compensatory changes in expression of other bicarbonate transporters (Slc4) are necessary to maintain calcification under acidic conditions [65]. Under extreme low pH conditions, TASK2 function may be under selection to ensure continued HCO3-uptake to regulate intracellular pH and facilitate skeletogenesis in sea urchin larvae. Empirical work is needed to validate this hypothesis.

**Figure 4:**
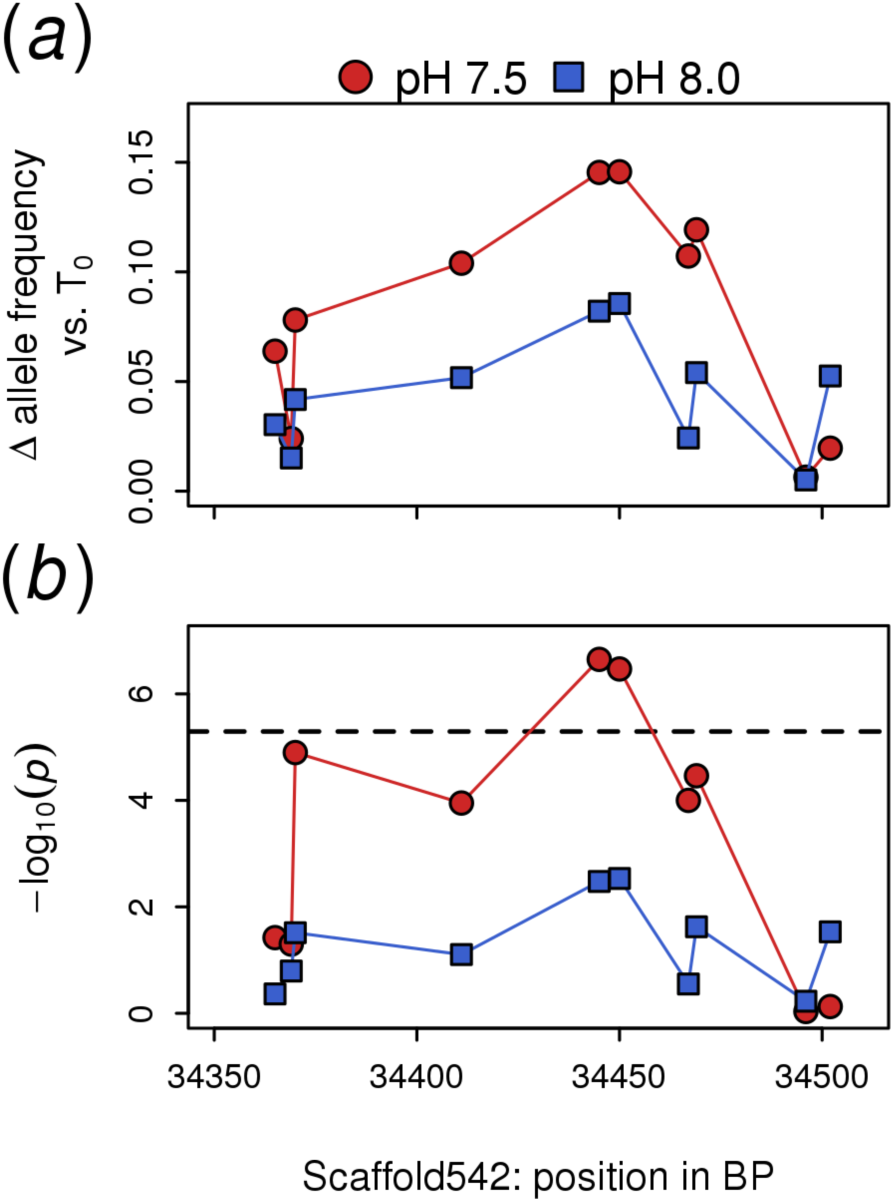
(a) Mean change in allele frequency of pH treatments relative to T_0_ across a ∼150 bp region of scaffold 542 in the K^+^ transporter TASK2. (b) *q*-values corresponding to the allele frequency changes in (a). The dashed horizontal line indicates the significance threshold of 5.09e-06.

A second protein of interest is a membrane bound and excreted enzyme in the carbonic anhydrase family (SPU_012518). Carbonic anhydrases catalyzes the hydration of CO_2_ to bicarbonate [66] and have been shown to be responsive to experimental acidification in many organisms including purple sea urchin [27,56], corals [67], anemones [68], mussels [69], oysters [70], and giant kelp [71]. In the present study, carbonic anhydrase (SPU_012518), appears to be responding to pH 7.5 where 3/12 variants have *P*-values < 0.01. However, these loci do not meet our stringent significance cutoff highlighting that results of any genome-wide scan for selection are sensitive to false negatives (and false positives), missing loci that may truly be under selection. However, given the strong signal of several loci across the gene, the established functional role in skeletogenesis in sea urchin larvae [72], and that the enzyme has been identified as pH-responsive in many ocean acidification studies across a broad range of taxa, carbonic anhydrase is an ideal candidate for future functional studies. Linking the genetic variation identified in the present study to functional phenotypes at the level of this enzyme would improve our understanding of the capacity of organisms to adapt to future conditions from standing genetic variation.

## Conclusion

We utilize a single generation selection experiment to reveal that purple sea urchins have genetic variation responsive to extreme and moderate low pH selection. The low variation among replicates, increased LD around selected loci, and enrichment for biological functions related to pH adaptation suggest that this approach accurately identifies adaptive loci. Further, we explore patterns of starting allele frequency to infer the evolutionary mechanisms underlying the maintenance of adaptive standing genetic variation. This work provides a framework for future studies assessing the genetic basis of adaptive responses to environmental stressors. Increasing the temporal sampling throughout development would reveal how different life stages differentially respond to selection, while functional studies of top candidates will help to validate their mechanistic and evolutionary significance. The results presented here show that *S. purpuratus* possesses genetic variation that is responsive to extreme low pH conditions and identifies a class of variants essential for adapting to low pH that are likely maintained by balancing selection. We also identify sets of rare variants that are unique to each pH selection regime suggesting that the preservation of large population sizes will be important in the maintenance of this rare adaptive variation.

## Data and code availability

Sequence data is available at the National Center for Biotechnology Information (NCBI SRA BioProject: PRJNA479817). Phenotype data is available as part of the supplementary material. All code for analysis is available at https://github.com/PespeniLab/urchin_single_gen_selection.

## Author Contributions

RB analyzed data and wrote the manuscript. AG designed and conducted the experiment and assisted with data analysis and drafting the manuscript. KH helped perform the experiment and analyzed the morphometric data. HH helped design aquaculture facilities and pilot the experiment. MP designed the experiment and wrote the manuscript.

## Acknowledgements

We thank Pete Halmay and Patrick Leahy for urchin collections. Jeremy Arenos for assistance with imaging and image analysis, Jason Hodin and Justin McAlister for helpful discussions on urchin larval culture, Lauren Ashlock for assistance with conducting the experiment, Mike Austin and the Vermont Advanced Computing Core for server space and maintenance, and all members of the Pespeni lab and Stephen Keller for helpful discussions.

