## Supplemental Methods and Figures for "Rare genetic variation and balanced polymorphisms are important for survival in global change conditions"

*Larval experiment:*

Purple sea urchin adults were collected in September 2016 from San Diego, CA, shipped overnight to the University of Vermont, and the experiment began immediately upon their arrival (n=25 total: 14 females and 11 males). For each female, 200,000 eggs were put into both moderate (pH 8.0) and extreme (pH 7.5) pH treatments (400,000 eggs per female). Twenty microliters of dry sperm from each male was pooled, diluted in 5 milliliters of ASW, and 100 microliters of this diluted sperm mix was added to each beaker of eggs for fertilization. After ensuring ≥ 95% fertilization success from each female, fertilized eggs were pooled by pH treatment and incubated at 14°C until the embryos hatched approximately 22 hours post-fertilization. Larval density was determined for each pool, and four replicate samples of 37,000 larvae were collected per pH, centrifuged to remove water, and flash frozen in liquid nitrogen for genomic DNA extractions. The remaining larvae were distributed into four replicate 3.7 L culturing vessels at each pH at a density of approximately 10 larvae $mL^{-1}$ (37,000 per replicate) and were cultured for 7 days post-fertilization. Larvae were fed green (*Dunaliella spp.*) and brown (*Isochrysis spp.*) algae, at 2500 cells/mL per algal species [1] on days 3 and 5 post-fertilization. Final genomic sampling of surviving larvae from the original 37,000 per replicate occurred day 7 post-fertilization in the same manner as day 1 sampling. An additional 12-33 larvae were preserved in calcium carbonate-buffered 10% formalin for morphometric analysis day 7 post-fertilization.

Larvae were cultured in recirculating systems designed in collaboration with Aqualogic (San Diego, CA). ASW, temperature, and pH were monitored and controlled in header tanks at 14°C and either 8.0 or 7.5, respectively, via a computer designed by RCK Controls (San Diego, CA) connected to Hamilton polilyte plus probes (Hamilton Company, Reno, NV). Instant Ocean ASW was used throughout the experiment at a salinity of 35ppt, although it is important to note that this artificial salt does not have the same total alkalinity (TA) as natural seawater. With a relatively high TA, the system was buffered too well and required high levels of $pCO_2$ to achieve the target pH levels (Table S1). However, pH levels were accurate. Therefore, this experiment likely consists of a stronger selective pressure than the pH alone due to high $pCO_2$ levels. While the water chemistry does not impact the objectives of this study, future experiments reliant on ASW should use one that more accurately mimics natural seawater chemistry conditions.

*Morphometrics*:

Larvae were photographed for morphometric analysis using a Photometrics Scientific CoolSNAP EZ camera (Tuscon, AZ) connected to a Zeiss Axioscop 2 compound microscope (Jena, Germany) at 10X. An image of a stage micrometer for size calibration was taken at 10X for each vessel at the time of imaging. Images were analyzed for total larval body length for 7-day old 4-armed pluteus larvae with ImageJ [2] by drawing a line that scaled the total length of the larva excluding arm lengths (where total length was defined as the distance from the base of the body to the oral hood, the bridge connecting the anterolateral arms), taking 3 replicate measurements, and averaging these replicates to obtain total body length per larva. The total number of larvae imaged in each replicate vessel varied from 12-33 larvae.

*DNA extractions & sequencing:*

DNA quality was assessed by running samples on an agarose gel and a NanoDrop while a Qubit 3.0 Fluorometer was used for quantification. DNA samples were sent to Rapid Genomics

(Gainesville, FL) for library preparation and capture-sequencing. For DNA library preparation, 250ng-1ug of genomic DNA from each sample was fragmented to an average size of 400 bp. DNA libraries were constructed by end-repair of the sheared DNA, A-tailing and adapter ligation, and barcoding followed by PCR amplification. DNA libraries were then captured with 46, 316 custom 120 bp probes designed by Rapid Genomics (Gainesville, FL) from the *S. purpuratus* genome (version 3.1, build 7). Probes were designed to capture exonic and regulatory regions of each gene in the genome (2 probes per gene), with regulatory probes being designed within 1000 bp upstream of the transcription start site, as determined by the *S. purpuratus* genome (version 3.1, build 7) gff3 file (http://www.echinobase.org/Echinobase/SpDownloads). The probes were hybridized to the DNA libraries and enriched for the specific targets. Samples were then pooled equimolar and sequenced using 100 bp paired-end reads on one lane of an Illumina HiSeq 3000.

*Mapping & SNP-calling:*

Raw paired-end reads were quality trimmed using Trimmomatic version 0.36 [3] with a phred score cut-off of 33, Illumina adapters removed, a sliding window value of 20:2, leading and trailing values of 2, and a requirement for reads to be a minimum length of 35 bp post-quality filtering and adapter trimming. Cleaned reads were then mapped via bwa mem [4] to the same genome with which the capture-sequencing probes were based upon (*S. purpuratus* genome 3.0 (build 7)), with a minimum seed length parameter of 5. Mapped reads were merged and sorted with Samtools version 1.2.1 [5] and sambamba version 0.6.0 [6] respectively, and indexed with samtools 1.2.1.

Using VCFtools version 0.1.15 [7], variant sites were filtered for minor allele frequency (maf) of 0.01, quality greater than 20, bi-allelic SNPs only, and no missing data. Because 25 parental individuals were used for crosses, we expect 0.02 (1 out of 50) to be the maf in these parental individuals. Therefore, a maf cutoff of 0.01 is reasonable. Sequencing depth can influence the accuracy of allele frequency estimation, and we use only variants where each pool was sequenced to a depth of > 40x and with an average minimum depth across all pools > 50x, as recommended by Schlotterer et al. [8]. Mean maximum coverage cutoff was 372. Finally, we removed off target variants, which were defined as any variant greater than 2kb from a probe. This filtering process resulted in 75,368 variant sites.

*Detecting changes in allele frequency:*

To test for increased LD after a single generation of selective mortality, we compared LD estimates among pairs of SNPs across the genome for selected and neutral variants using LDx [10] where a selected locus is defined as having a CMH q-value < 5.09e-06. Reads were considered with a minimum read depth of 40, maximum depth of 4800, distance between read pairs of 500, quality score of at least 20, minor allele frequency of 0.05, and a minimum intersection depth of 10 (*-l 40 -h 4800 -s 500 -q 20 -a 0.05 -i 10*). LD decay was assessed by fitting an exponential decay curve using the R package *nlme* [11]. Note that LD estimates were only used for variants within 100 base pairs (bp) as estimates become noisy due to the low number of variants greater than this distance. Altering this cutoff to 150 (bp) results in the same general LD patterns seen in Fig. 3. Permutation tests were run as described in the main text.

A permutation test was run to check for bias in the CMH test statistic towards identification of alleles starting at low frequency as significant. The expected change in allele frequency in response to selection is proportional to the product of the major and minor allele frequency and it is expected that the CMH statistic should be biased against detecting shifts in low frequency alleles [12]. Here, we randomly assigned labels to each of the 12 samples (four replicates from each of $T_0$, pH 7.5, pH 8.0) and calculated the CMH test statistic for each locus for these permuted samples and repeated this 500 times. Given the random treatment assignment, any CMH signal is a false positive. Therefore, if the test statistic is biased towards identifying loci with specific allele frequencies, this permutation should reflect that. For each permutation, the allele frequencies of the loci from the top 0.015 quantile of p-values were compared against our observed allele frequency distributions using Wilcoxon rank-sum tests.

*Quantifying strength of selection*

To determine the relative strength of selection in each treatment, we integrated larval density estimates with effective population size and estimates of the selection coefficient. Larval densities were estimated at 7 days post-fertilization using each of the four replicates per pH treatment. 10mL samples were taken from each culture vessel and added to 5mL of 10% formalin (3.3% final concentration). Samples were kept at 4C for 2-3 days, allowing the preserved larvae to settle at the bottom. To count the 7-day old larvae to determine final density, samples were centrifuged, all but 800 ul of liquid was removed, and larvae were mounted onto a slide and counted under a compound microscope. Final density (larvae/mL) was then multiplied by the volume in each vessel (3,700 mL) to determine the final number of larvae that survived. Finally, we define the realized selection coefficient (S) following Kessner and Novembre [13] as the selection coefficient that would result in the change in allele frequency given a specific allele frequency. We calculate this as $\Delta p/((p*(1-p))-(2*p*\Delta p))$ and compare genome wide estimates using a Wilcoxon rank-sum test.

**Supplemental Tables:**

**Table S1:** Mean water chemistry measurements during the selection experiment

| Treatment | Temp (°C) | Salinity (ppt) | pH | TA (µmol*kg$^{-1}$) | pCO2 (µatm) |
|---|---|---|---|---|---|
| **8.0** | 14.39 $\pm$ 0.14 | 33.04 $\pm$ 0.16 | 8.04 $\pm$ 0.05 | 3460 $\pm$ 8.11 | 703.04 $\pm$ 48.89 |
| **7.5** | 14.48 $\pm$ 0.14 | 33.53 $\pm$ 0.25 | 7.59 $\pm$ 0.04 | 3545 $\pm$ 8.63 | 1928.89 $\pm$ 39.24 |

**Table S2:** Detailed information for all variants. Information includes allele frequency estimates for each treatment, mean and standard deviation of change in allele frequency, and gene annotation.

**Table S3:** GO enrichment results. "set" indicates the pH treatment for each term.

**Table S4:** Morphometrics data from Figure 1.

**Supplemental Figures:**

**Figure S1:** Manhattan plot of the CMH Q-value for (A) pH 7.5 and (B) 8.0. Alternating black and grey points are alternating scaffolds of the *S. purpuratus* reference genome. Significant variants are indicated by red circles (pH 7.5), blue squares (pH 8.0), or purple triangles (significant in both treatments).

**Figure S2:** Violin plot of changes in allele frequency. Horizonal line in each plot indicates the median value.

Figure S3: Allele frequency at $T_0$ where frequencies are polarized by the rising allele. Colored solid lines are the density plot distributions for each set of selected loci. The solid black line represents the median expectation of neutral loci from 1000 permutations of down sampling to the same number of variants as selected loci; grey shading is the 95% distribution of the permutation. Dashed vertical lines are the mean allele frequencies of each set of variants.

**Figure S4:** Results from CMH permutation test. Sample labels were shuffled and allele frequencies of the top 0.015 quantile of p-values were considered "significant". The solid black line is the median of the permutation allele frequency distribution with the 95% distribution shown as grey shading.

**Figure S5:** Change in allele frequency versus folded minor allele frequency.

**Figure S6:** Relative frequencies of SNP classes in response to selection.

**Figure S1**

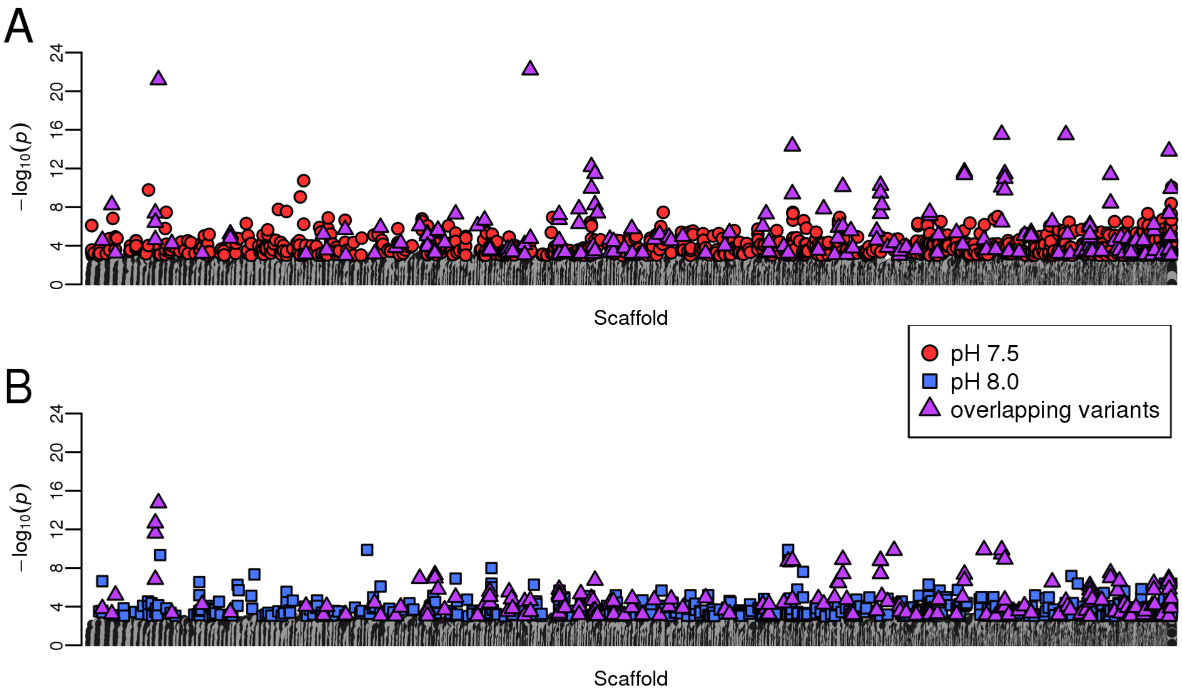


**Figure S2**

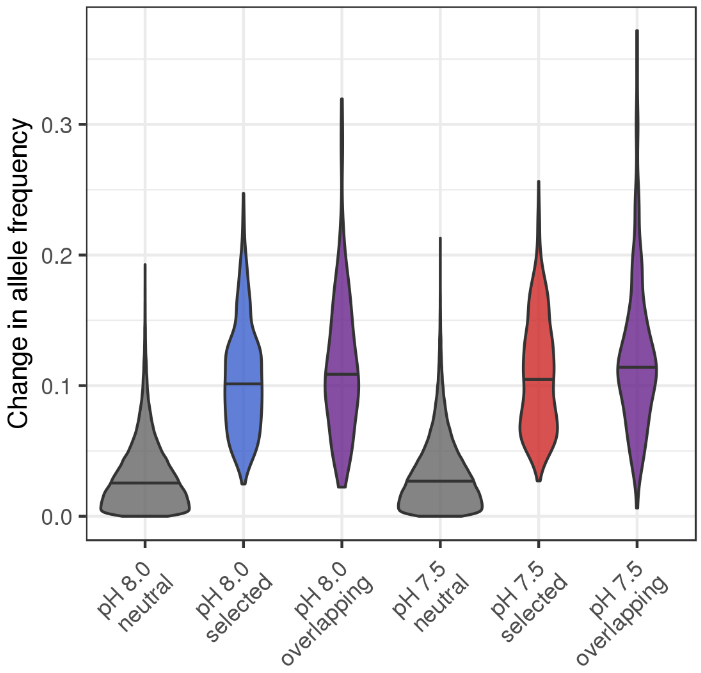

**Figure S3**

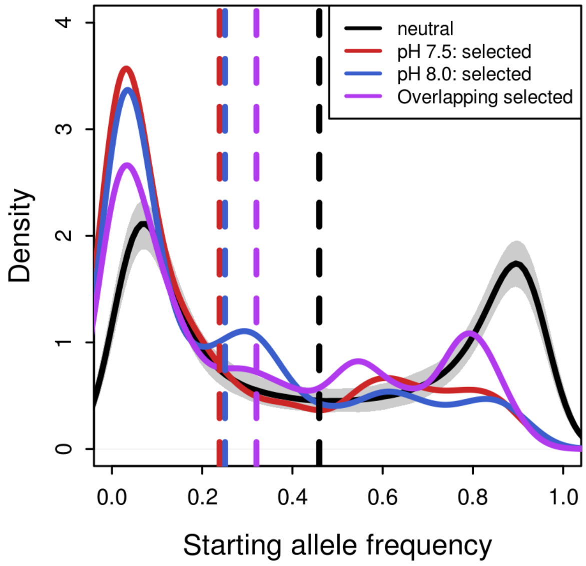


**Figure S4**

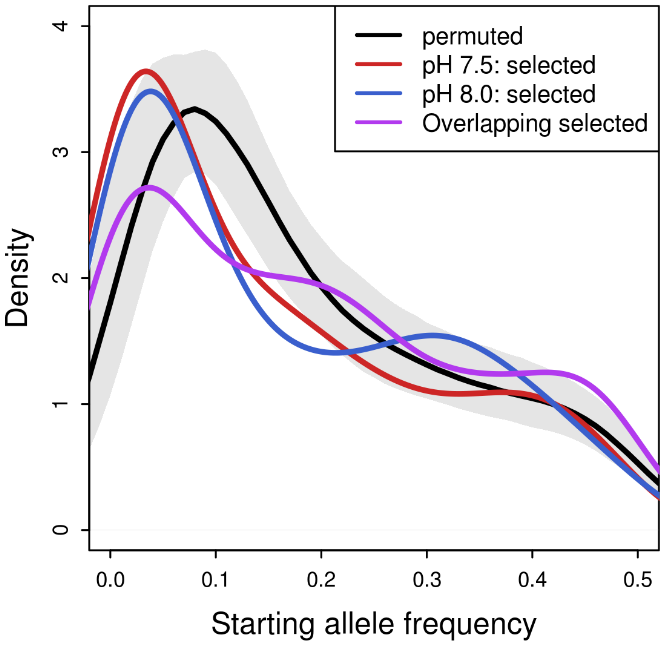

### Figure S5

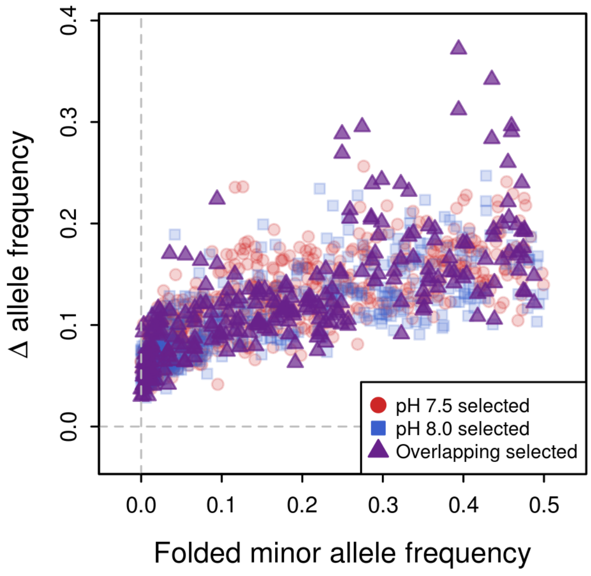

0.4
0.3
0.2
0.1
0.0
Δ allele frequency
0.0
0.1
0.2
0.3
0.4
0.5
Folded minor allele frequency
pH 7.5 selected
pH 8.0 selected
Overlapping selected

**Figure S6**

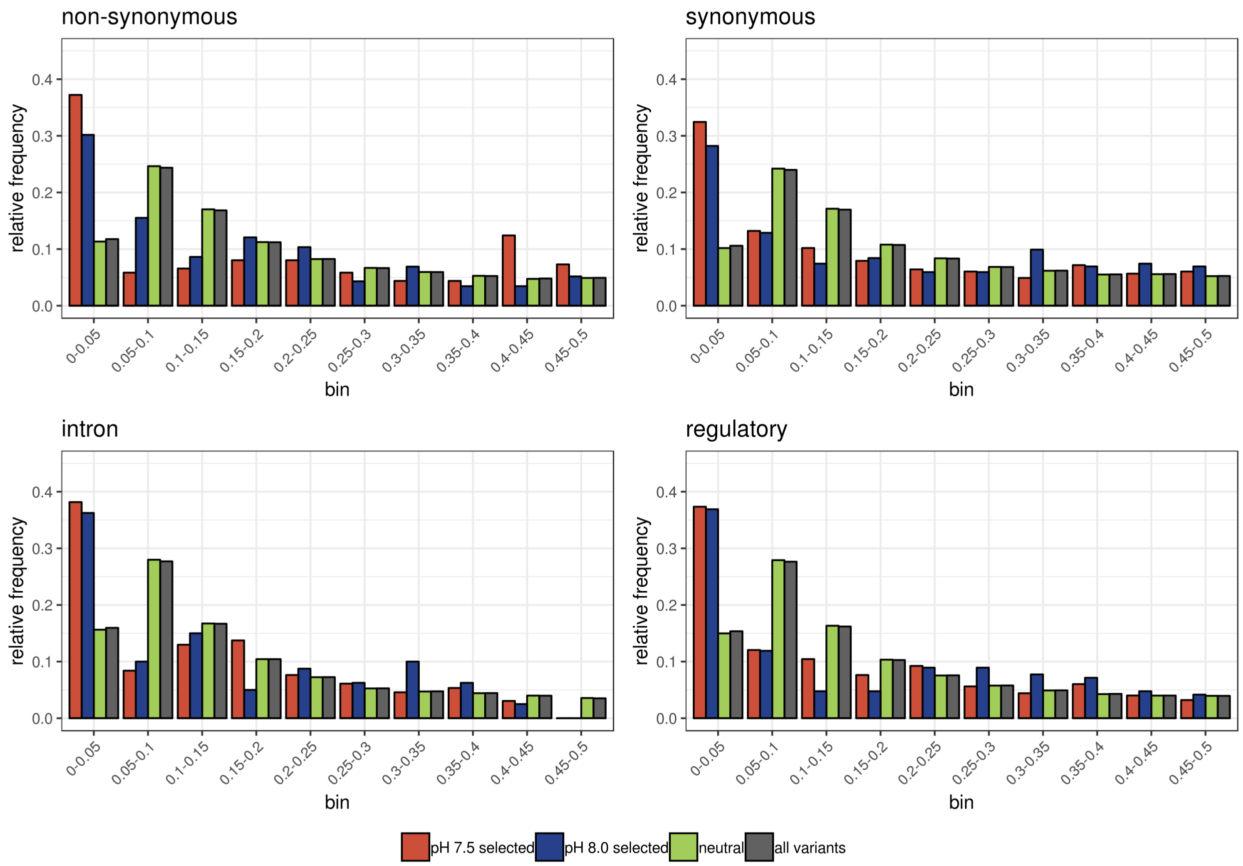

non-synonymous
synonymous
intron
regulatory
relative frequency
0.0
0.1
0.2
0.3
0.4
0-0.05
0.05-0.1
0.1-0.15
0.15-0.2
0.2-0.25
0.25-0.3
0.3-0.35
0.35-0.4
0.4-0.45
0.45-0.5
bin
pH 7.5 selected
pH 8.0 selected
neutral
all variants
